# Cryo-EM structures of both ends of the actin filament explain why the barbed end elongates faster than the pointed end

**DOI:** 10.1101/2023.05.12.540494

**Authors:** Steven Z. Chou, Thomas D. Pollard

## Abstract

Actin filament ends are the sites of subunit addition during elongation and subunit loss during depolymerization. Prior work established the kinetics and thermodynamics of the assembly reactions at both ends but not the structural basis of their differences. Cryo-EM reconstructions of the barbed end at 3.1 Å resolution and the pointed end at 3.5 Å reveal distinct conformations at the two ends. These conformations explain why barbed ends elongate faster than pointed ends and why pointed ends rapidly dissociate the γ-phosphate released from ATP hydrolysis during assembly. The D-loop of the penultimate subunit at the pointed end is folded onto the terminal subunit, precluding its binding incoming actin monomers, and gates on the phosphate release channels of both subunits are wide open. The samples were prepared with FH2 dimers from fission yeast formin Cdc12. The barbed end reconstruction has extra density that may be partial occupancy by the FH2 domains.

**Significance Statement:** Cells depend cytoplasmic filaments assembled from the protein actin for their physical integrity, as tracks for myosin motor proteins and movements of the whole cell and internal organelles. Actin filaments elongate and shrink at their ends by adding or dissociating single actin molecules. We used cryo-electron microscopy to determine the structures of the two ends of actin filaments at 3.5 Å resolution for the slowly growing pointed end and 3.1 Å for the rapidly growing barbed end. These structures reveal why barbed ends grow faster than the pointed ends, why the rate at the pointed end is not diffusion-limited and why the pointed end has a low affinity for the γ-phosphate released from bound ATP inside the filament.

## Introduction

Assembly and disassembly of actin filaments takes place exclusively by the addition and loss of actin subunits at the two ends making detailed knowledge about the structures of the two ends essential for understanding the mechanisms. For decades, without thinking about it too seriously, the field assumed that the structures of the ends are similar to internal subunits, so new subunits would associate with the ends by forming bonds similar to those inside the polymer. Since the mid-1970s a series of discoveries revealed that the two ends must differ from each other and the internal subunits. High resolution structures in this paper show how the two ends differ.

The first evidence that the ends differ came from electron microscopy showing that the barbed end (defined by the polarity of myosin heads bound to the filament) grows faster than the pointed end (Woodrum, Rich et al. 1975); (Hayashi and Ip 1976); (Kondo and Ishiwata 1976)). Measurements of the rate constants for association and dissociation of subunits at the two ends for ATP- and ADP-actin monomers explained quantitatively why the rates differ (Pollard and Mooseker 1981); (Pollard 1986)). The dependence of elongation rate constants on solution viscosity revealed that elongation at the barbed end is a diffusion-limited reaction with a high orientation factor of 0.02 (binding occurs during 2% of collisions between monomers and the barbed end), while elongation at the pointed end is not diffusion-limited (Drenckhahn and Pollard 1986). Kinetic analysis also revealed that the equilibrium constants at the two ends are the same for ADP-actin (as expected) but differ during elongation by ATP-actin monomers (Pollard 1986). This difference arises from different affinities of the two ends for phosphate (Fujiwara, Vavylonis et al. 2007) and explains the slow flux (treadmilling) of subunits through filaments at steady state in the presence of ATP-actin monomers (Wegner 1976); (Pollard 1986).

Using x-ray fiber diffraction Oda et al. (Oda, Iwasa et al. 2009) discovered that the subunits in actin filaments are flattened compared with monomers. Subsequent higher resolution cryo-EM structures (Fujii, Iwane et al. 2010); (Merino, Pospich et al. 2018); (Chou and Pollard 2019); (Oda, Takeda et al. 2019) established that flattening occurs during incorporation of subunits into the filament rather than later during ATP hydrolysis and phosphate release, since the subunits are flattened in AMPPNP-, ADP-P_i_- and ADP-actin filaments (Chou and Pollard 2019). A 23 Å resolution cryo-EM structure showed that the conformations of the last two subunits at the pointed end of the filament differ from internal subunits (Narita, Oda et al. 2011) with the penultimate subunit bent against the terminal subunit, interfering with the addition of new subunits. No differences between the internal and terminal subunits could be distinguished in a 23 Å resolution cryo-EM structure of the barbed end bound to capping protein (Narita, Takeda et al. 2006).

During all atom molecular dynamics (MD) simulations of short filaments cut from longer ATP-, ADP-P_i_- or ADP-actin filaments (Zsolnay, Katkar et al. 2020), both ends relaxed to new conformations during a few hundred nanoseconds of computed time. The terminal subunits at both ends were more twisted than the subunits in their parent filaments, but the internal subunits flattened as they acquired new neighbors. The new pointed end spontaneously adopted a structure similar to the Narita structure (Narita, Oda et al. 2011) with the residues in the D-loop of the penultimate subunit P-1 only 5% as accessible to the solvent as those of an actin monomer. Most remarkably, terminal subunit B at the barbed end lost its lateral bonds with subunit B-1, so it was tethered loosely by its D-loop to subunit B-2. This is reasonable, since neither subunit B nor subunit B-1 are flattened, which compromises their lateral interactions (Chou and Pollard 2019).

Here we present high resolution cryo-EM structures of both ends of the actin filament, which explain five decades of observations on the different elongation rates and affinities for phosphate at the two ends.

## Results

### Quality of the EM structures

Imaging enough ends for high resolution reconstructions has been an historic challenge for the field given the long lengths typical of actin filaments, but the FH2 domains of formin Cdc12 from fission yeast made this possible, because they nucleate filaments that grow very slowly at both ends (Kovar, Kuhn et al. 2003). By optimizing cryo-EM vitrification conditions, we generated numerous short filaments, a few with obvious FH2 caps but most appearing bare (Fig. S1). Loss of the Cdc12 FH2 domains was unexpected, since they are highly processive (Kovar, Harris et al. 2006). One possibility is that some FH2 domains dissociated during interactions with carbon support film. By optimizing data acquisition, we collected images of ∼450,000 short filaments with both ends available for 3D reconstructions (Table 1). After particle picking, classification and reconstruction, the resolutions were 3.1 Å at the barbed end and 3.5 Å at the pointed end (Fig 1, Table 1).

**Table 1.** Statistics for data collection, map reconstruction and model refinement

|  | Pointed end<br>(EMD-*****, PDB *****) | Barbed end [with B subunit]<br>(EMD- *****, PDB *****) |
| --- | --- | --- |
| **Data collection and map reconstruction** |  |  |
| Magnification | 46816 |  |
| Voltage (kV) | 300 |  |
| Electron exposure (e<sup>-</sup>/Å<sup>2</sup>) | 56.56 |  |
| Defocus range (μm) (set/measured) | (-2.5, -1.5)/(-3.5, -1.0) |  |
| Pixel size (Å) | 1.068 |  |
| Symmetry | C1 |  |
| Initial particle images (no.) | 448222 | 506236 |
| Final particle images (no.) | 448222 | 471000 |
| Map analysis in reconstructed/modeled regions |  |  |
| Resolution (Å) at 0.143 threshold, Relion | 3.5 | 3.1 [3.2] |
| Sharpening B factor (Å<sup>2</sup>), Relion | -114.94 | -114.51 [-132.7] |
| Map resolution range (Å) | 3.1-4.3 | 2.9-4.1 [2.9-5.9] |
| **Model Refinement** |  |  |
| Initial model used (PDB ID) | 6DJN | 6DJN |
| Model-to-map fit correlation | 0.74 | 0.68 [0.67] |
| Model resolution (Å) at 0.5 threshold | 3.5 | 3.6 [3.8] |
| Model composition |  |  |
| Heavy atoms | 17356 | 17632 [20574] |
| Protein residues | 2196 | 2232 [2604] |
| Ligands (ATP/ADP/Pi/Mg<sup>2+</sup>) | 0/6/0/6 | 0/6/2/6 [1/6/2/7] |
| B factors (Å<sup>2</sup>) |  |  |
| Protein | 99.72 | 80.64 [108.89] |
| Ligands | 92.95 | 73.40 [106.44] |
| R.m.s. deviations |  |  |
| Bond lengths (Å) | 0.01 | 0.01 [0.01] |
| Bond angles (°) | 1.43 | 1.24 [1.24] |
| Geometry validation |  |  |
| MolProbity score | 1.87 | 1.79 [1.78] |
| Clashscore | 7.09 | 6.29 [6.40] |
| rotamer/Cβ outliers (%) | 0.27/0.00 | 0.21/0.00 [0.18/0.00] |
| Ramachandran plot |  |  |
| Favored (%) | 92.32 | 93.05 [93.57] |
| Allowed (%) | 7.68 | 6.95 [6.43] |
| Disallowed (%) | 0.00 | 0.00 [0.00] |

**Figure 1.**
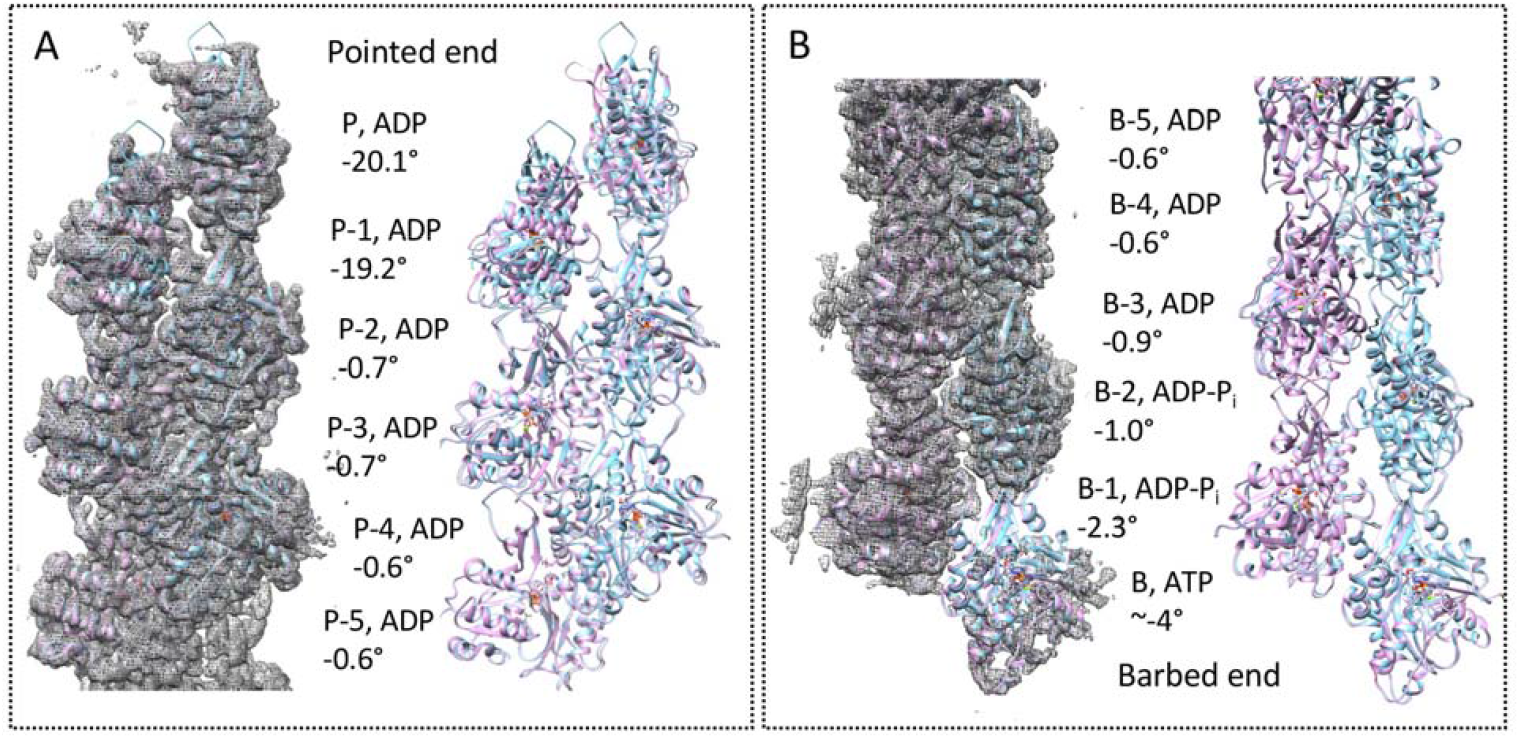
Overview of the structures of the pointed and barbed ends of actin filaments with the bound nucleotides and dihedral angles between subdomains-1/2 and subdomains-3/4 of each subunit. A, pointed end; B, barbed end. Each panel shows the map of the reconstruction from cryo-electron micrographs and ribbon diagrams of two models: plumb color, model of the end; and sky-blue color, model of a standard ADP-P_i_-actin filament (PDB 6DJN) fit to internal subunits and extended to the terminal subunit. The subunits at the pointed end are named P, P-1, P-2, P-3, P-4 and P-5. The subunits at the barbed end are named B, B-1, B-2, B-3, B-4 and B-5. The dihedral angles between subdomains 1-2 and subdomains 3-4 are given in degrees.

The high quality of the maps allows identification of the nucleotides bound to all 13 actin subunits (Figure 2). All internal subunits in both structures have bound ADP. Subunits P and P-1 at the pointed end also have bound ADP. At the barbed end subunit B has bound ATP, while subunit B-1 and B-2 have bound ADP with additional densities for a separated γ-phosphate about 30% as strong as the α- and β-phosphates.

**Figure 2.**
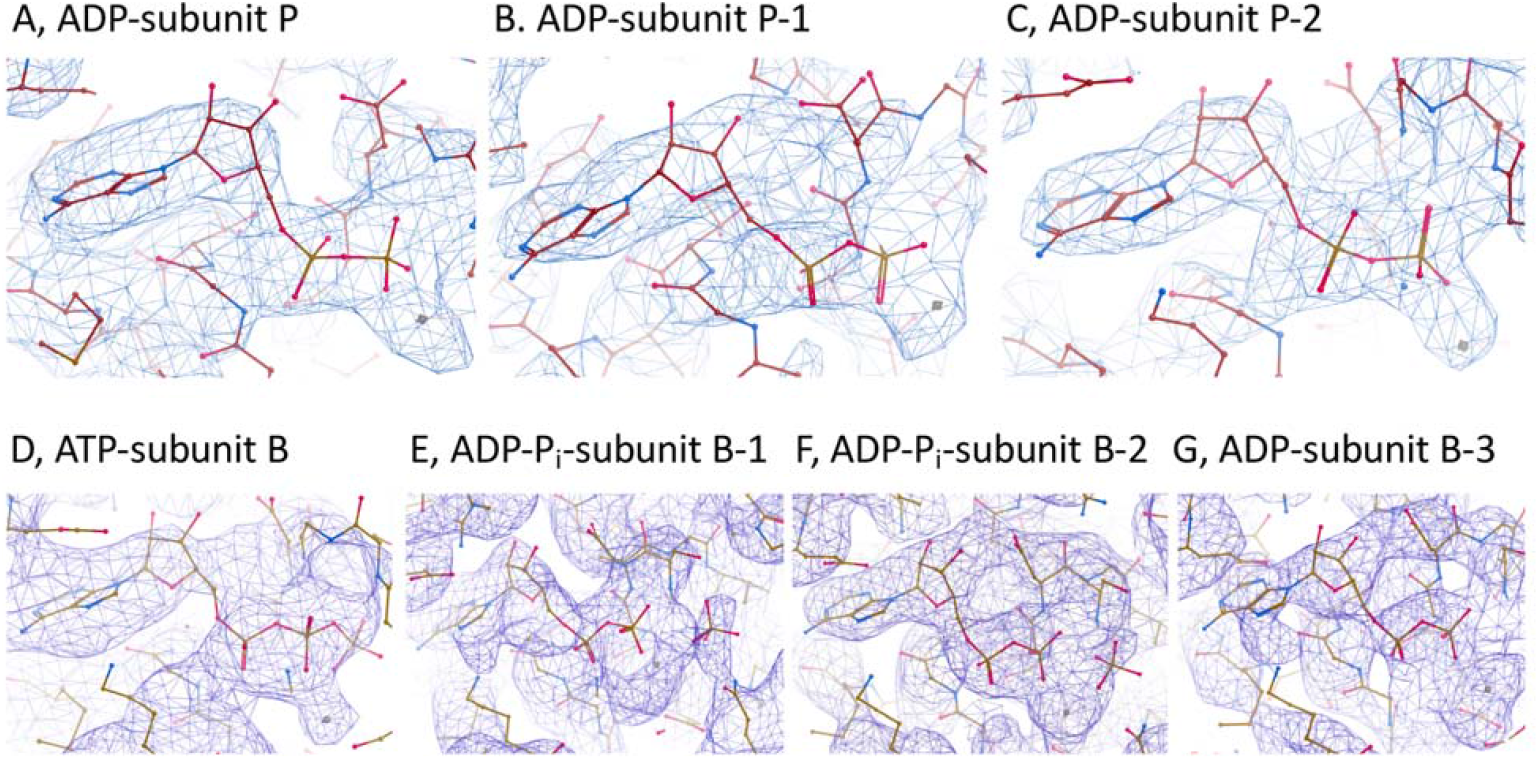
Map densities (blue) from the terminal subunits at both ends fit with stick diagrams (red) of models of the bound nucleotides and parts of the surrounding actin subunits. Magnesium ions are grey spheres.

The reconstructions also offer insights into the release of the γ-phosphate cleaved from ATP during assembly. The side chain of R177 in subdomain 3 forms a gate on the exit channel (Wriggers and Schulten 1997), which closes when R177 forms a hydrogen bond about 3 Å long with N111 in subdomain 1 in AMPPNP- and ADP-P_i_-actin filaments (Chou and Pollard 2019). This gate is open with ∼10.0 Å separating N111 and R177 in Ca-ATP-actin monomers (PDB 2A42) and Mg-ADP-actin monomers (PDB 3A5L), and slightly open in one (Chou and Pollard 2019) and closed in other reconstructions of ADP-actin filaments (Oosterheert, Klink et al. 2022, Reynolds, Hachicho et al. 2022). Judging from the distances between the side chains of N111 and R177, the backdoors on the phosphate release channel are wide open in subunits P (9.2 Å) and P-1 (8.2 Å), slightly open in subunits B (∼5.3 Å) and B-1 (4.1 Å) and closed in all other subunits at both ends (Fig. 3 and Table 2). The side chains of both N111 and R177 rotate toward each other to close the gate.

**Table 2.**
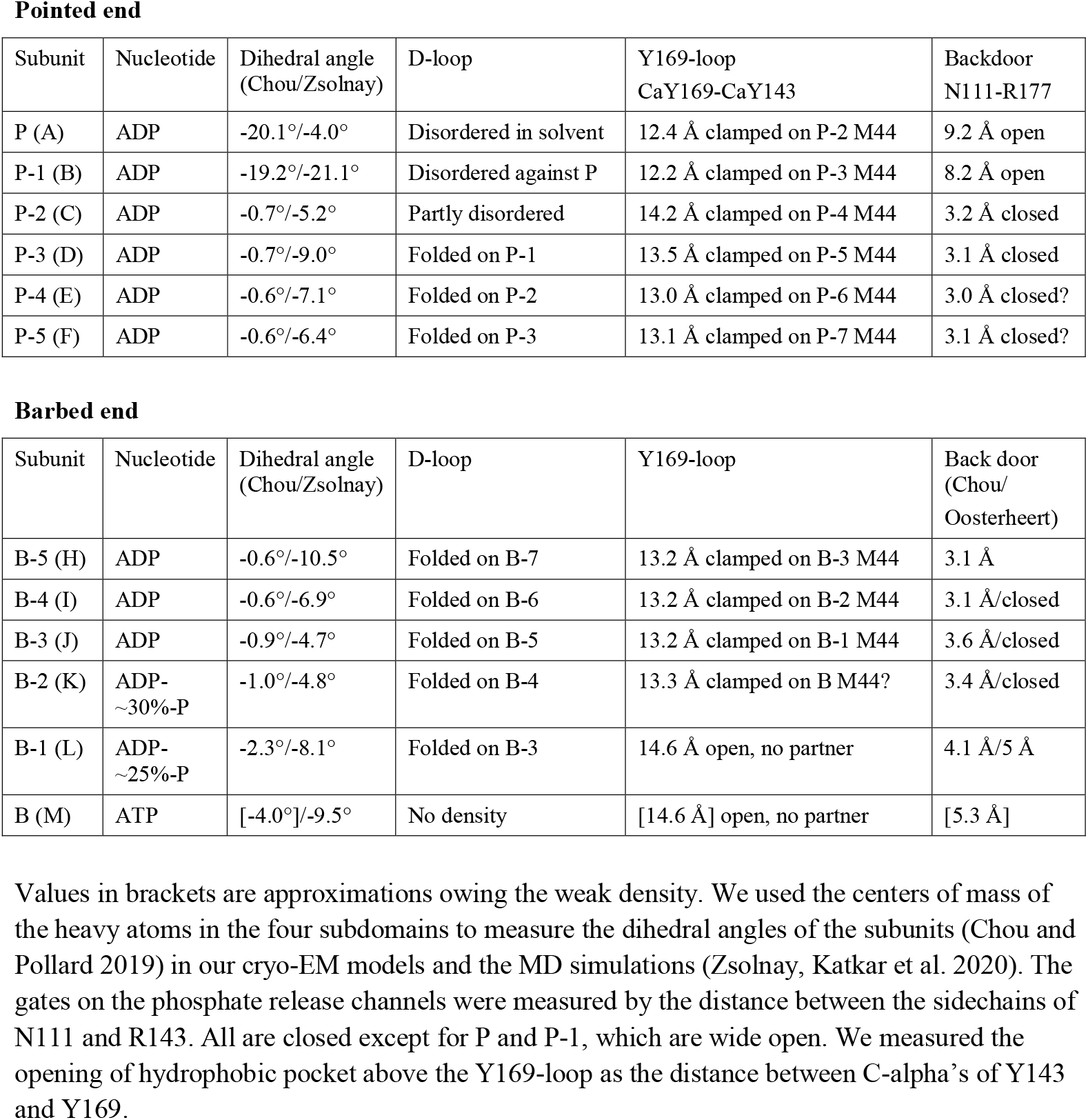
Summary of structural data with comparisons to Zsolnay et al. (Zsolnay, Katkar et al. 2020) and Oosterheert et al. (Oosterheert, F.E.C. et al. 2023).

**Figure 3.**
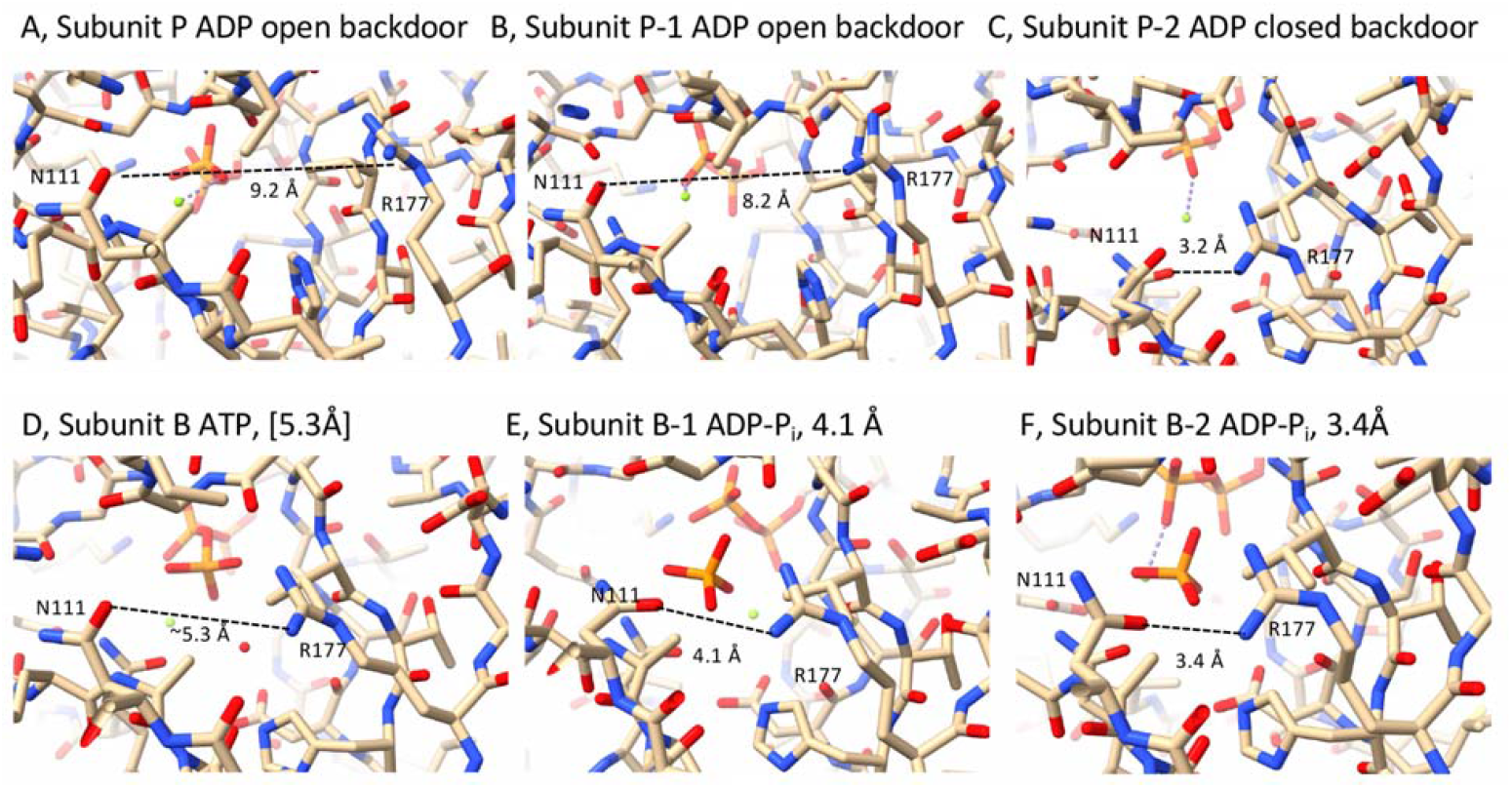
Comparison of the conformations of the backdoor residues of subunits at both ends of the filament illustrated with stick figures. The dashed black lines show the distances between the sidechains of residues N111 and R177, which form the gate on the phosphate release channel. These gates are open in subunits P and P-1, slightly open in subunits B and B-1 and closed in all other subunits at the two ends. Panels A-C show the β-phosphate in orange and red. Panels D-F show the γ-phosphate in orange and red behind the backdoor.

### Structure of the pointed end

Our high-resolution model of the subunits of ADP-actin filaments (Chou and Pollard 2019) fit well into map of the internal part of the pointed end reconstruction (Fig 1A) but not the terminal two subunits. As expected from a 22.9 Å reconstruction (Narita, 2011) and MD simulations of the pointed end (Zsolnay, Katkar et al. 2020), the conformations of the two terminal subunits differ from the internal subunits (Figs. 1, 3 and 4).

**Figure 4.**
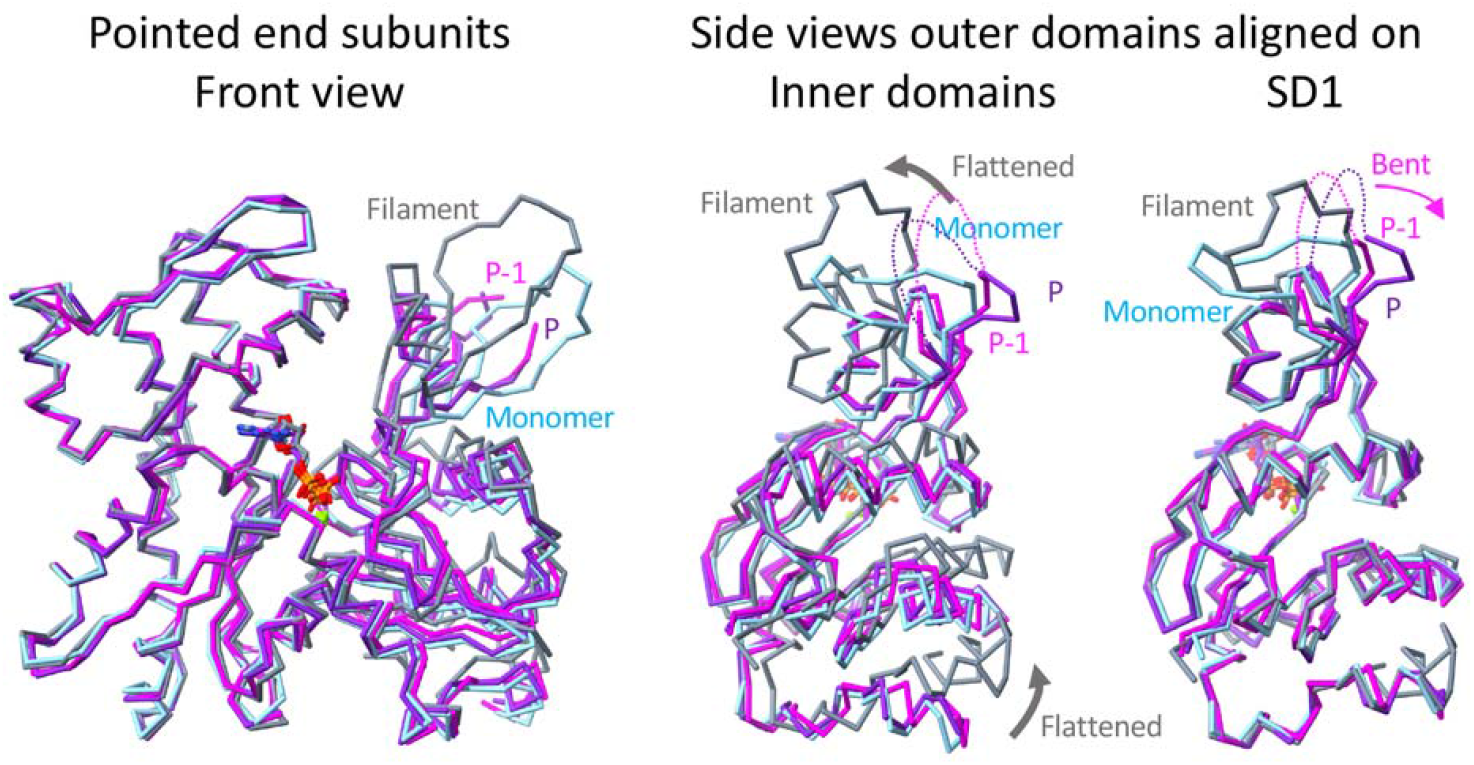
Comparisons of the conformations of the terminal subunits at the pointed end by superimposing backbone traces of a typical internal subunit (B-6, grey), an actin monomer (sky-blue; PDB 2A42), subunit P (magenta) and subunit P-1 (purple). The disordered D-loops of subunits P and P-1 are missing the front view and shown as dotted lines in the side views. The grey arrows in the middle panel indicate the flattening of the internal subunit (grey) relative to the monomer (sky-blue).

#### Terminal subunit P

With a dihedral angle of -20.1°, subunit P is more twisted than an actin monomer, because SD2 is bent backwards relative to SD1 (Fig. 4 right) by pivoting around five points (loop with L94, loop before T103, loop with E107, loop with L110, and the loop after helix N137 to Y143) (Fig. 4). The map has no density for the disordered D loop between G42 and K50, which judging from the breaks in the backbone, points into the solvent . The backdoor for phosphate release is open, because the side chain of R177 twists inward (Fig. 3A) rather than forming the hydrogen bond with N111 that closes the channel (Fig. 3C). The open door explains why phosphate is absent from the active site. The Y169 loop (also known as the W-loop) is clamped around the side chain of subunit P-2 M44. At the C-terminus of subunit P, the final helix is tilted toward the barbed end and density for the backbone ends at A365.

#### Penultimate subunit P-1

The overall conformation is similar to that of subunit P with most of subdomains 1, 3 and 4 in the standard actin filament conformation), but subdomain 2 is bent backwards giving a dihedral angle of -19.2° between subdomains 1-2 and subdomains 3-4 (Fig 4). The map has no density for disordered D loop between Q41 and K50, which judging from the breaks in the backbone, points toward the hydrophobic loop residues A319, L320, A321 and P322 of subunit P, as observed in molecular dynamics simulations (Zsolnay, Katkar et al. 2020). Fig. 5D shows interactions of the D-loop with the hydrophobic plug at two time points during an MD simulation. The backdoor for phosphate release is open similar to subunit P (Fig. 3B). The Y169 loop is clamped around the side chain of subunit P-3 M44. The density for the backbone also ends at A365.

**Figure 5.**
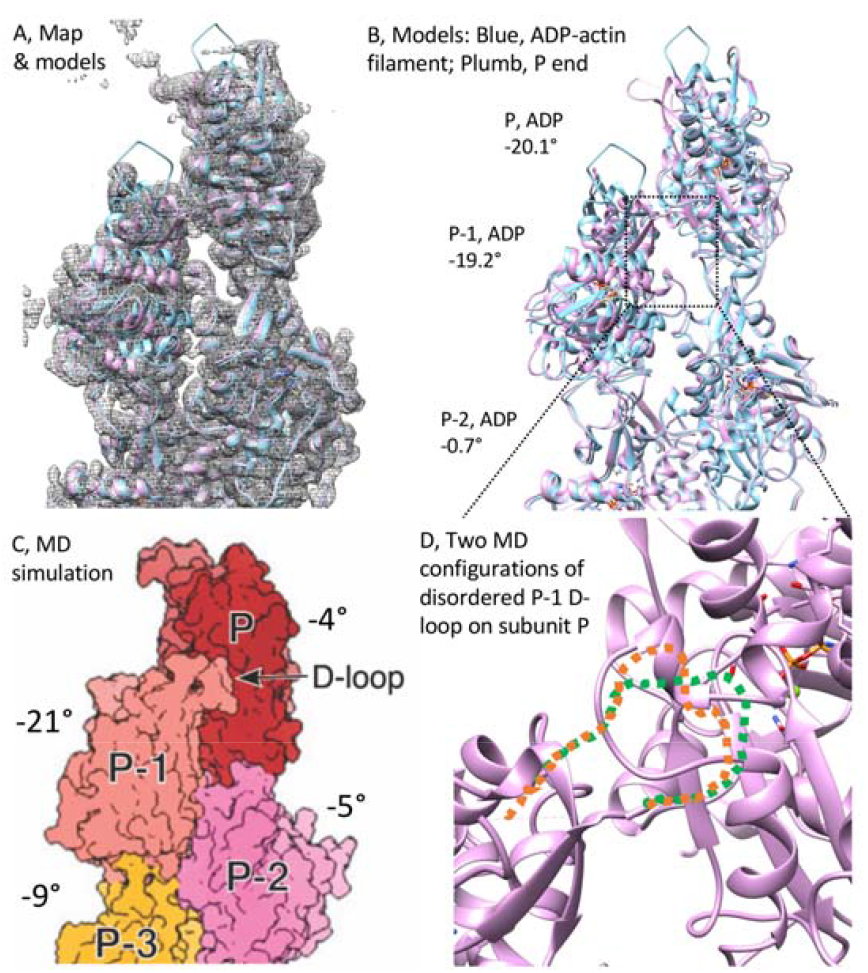
Details of the pointed end structure with bound nucleotides and dihedral angles in degrees. A. Map with two models of the three terminal subunits. B, Ribbon diagrams of two models; plumb color, new model of the end fit into the map; and sky-blue color, structure of a standard ADP-P_i_-actin filament (PDB 6DJN) fit to internal subunits and extended to the terminal subunit. C, Model of pointed end from the MD simulations of Zsolnay et al. (Zsolnay, Katkar et al. 2020) relabeled with dihedral angles calculated from the centers of mass of the heavy atoms in the four subdomains. D, Ribbon diagram showing the lateral interaction of subunit P-1 with the hydrophobic plug on the side of subunit P. The map has no density for the disordered D-loop of subunit P-1, which assumes a variety of positions on the adjacent hydrophobic plug of subunit P in MD simulations. The green and orange dashed lines show positions of the D-loop at two points in time from MD simulations.

#### Subunit P-2

The conformation is very similar to the rest of the internal subunits but P-2 has no density for D-loop residues V45, G46 M47 and G48 in spite of the side chain of side chain of its residue M44 being clamped in the hydrophobic pocket above the Y169-loop of subunit P. This pocket in subunit P is open more widely than standard actin filament subunit.

### Structure of the barbed end

Our 3.1 Å resolution reconstruction of the barbed end (Figs. 1B and 6) shows that subunits internal to subunit B-2 are flattened (Table 2) and have bound ADP (Fig. 2G) like our model of the ADP-actin filament (Chou and Pollard 2019). The filament model fits less well into the maps of the 3 terminal subunits (Figs. 1, 4 and 6), which are slightly twisted but flatten progressively toward the interior (Table 2).

**Figure 6.**
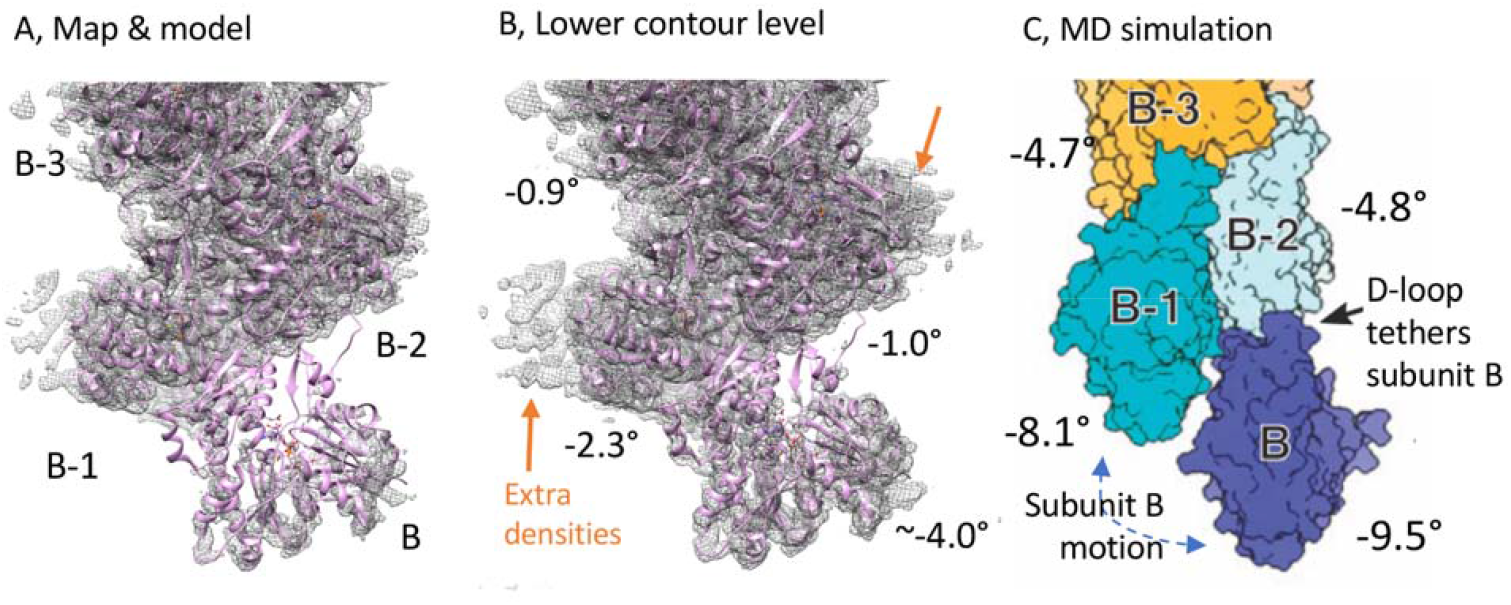
The barbed end structure with dihedral angles in degrees. A, Map at a standard contour level fit with a ribbon diagram of the new model. The density for the terminal subunit B is low. B, Map with a lower contour level with a more complete density for subunit B and the extra densities associated with the sides of subunits B-1 and B-2. C, Model of pointed end from the MD simulations of Zsolnay et al. (Zsolnay, Katkar et al. 2020) relabeled with dihedral angles calculated from the centers of mass of the heavy atoms in the four subdomains.

#### Terminal subunit B

The map has low density for a subunit at the barbed end of the filament, as anticipated by MD simulations showing the terminal subunit flexibly tethered by its D-loop to the subunit B-2 (Zsolnay, Katkar et al. 2020). We interpret this density as the terminal subunit B. Note that both ends were processed the same way, but only the barbed end has this weak density for a terminal subunit. The map of subunit B has convincing density in the active site for Mg-ATP (Fig. 2D) and helices in SD1 and SD3 but not SD4. The map lacks density for most of SD2 including the D-loop, but the hydrophobic cavity above the W-loop of subunit B-2 is partially open as if clamped on the side chain of M44 from the D-loop of subunit B. The backdoor gate of the phosphate release channel may be slightly open (∼5.3 Å), although this distance is an estimate given the low density of the map.

#### Subunits B-1 and B-2

The maps of subunits B-1 and B-2 both have strong densities for bound ADP and lower densities for separated γ-phosphates in the active site (Fig. 2EF). The gate of the phosphate release channel of subunit B-1 may be partially open as the distance between the side chains of N111 and R177 (4.1 Å) is slightly larger than internal subunits (3.1-3.6 Å, Table 2).

### Extra densities at the barbed end

Contouring the map of the barbed end at lower levels emphasizes weak extra densities associated with the sides of subunits B-1 and B-2 and to a lesser extent with B-3 (Fig. 6). The reconstructions of both ends were processed the same way, but these peripheral densities appear only at the barbed end not the pointed end (Fig. 5). Formin FH2 domains might contribute to these densities. These densities were present in the barbed end reconstruction in spite of the fact that during classification of those ends, we removed about 5% of the particles which appeared to have extra, peripheral densities and processed them separately. The resolution of this subset was limited by the small number of particles, preferred orientation and possibly by heterogeneity of the extra densities.

## Discussion

### Pointed end structure

Our cryo-EM reconstruction of the pointed end shows that the two terminal subunits, P and P-1, have twisted conformations with dihedral angles of -20.1° and - 19.2° more like monomers than the flat internal subunits in the same filament. However, the twisting is largely between subdomains 1 and 2 of these subunits rather than between subdomains 1-2 and subdomains 3-4 as in actin monomers (Fig. 4). In addition, subdomain 2 of subunit P-1 bends over, allowing the D-loop to contact the side of subunit P (Fig. 5D).

Our cryo-EM reconstruction confirms at high resolution features observed in previous studies. The pioneering 23 Å reconstruction of Narita et al. (Narita, Oda et al. 2011) was the first look at the distorted confirmations of the two terminal subunits, including tilting of part of subunit P-1 onto terminal subunit P. Although their map did not allow tracing the polypeptide backbone, the shifts in mass led to their accurate interpretation that the D-loop of subunit P-1 interacts with the hydrophobic plug of terminal subunit P. We agree with their interpretation that this conformation would inhibit both subunit association and dissociation at the pointed end.

Zsolnay et al. (Zsolnay, Katkar et al. 2020) used all atom molecular dynamics simulations to document conformational changes after creating a new pointed end in standard AMPPNP-, ADP-P_i_- and ADP-actin filaments (Chou and Pollard 2019). During the first 200 ns of simulated time, the two terminal subunits at new pointed end of ADP-actin filaments relaxed from the flattened conformation of internal subunits to a more twisted conformations. In the simulations, the flexible D-loop of subunit P-1 explored the hydrophobic surface of the adjacent hydrophobic plug of subunit P (Fig. 5D). Our map lacks density for the D-loop, so we use two snapshots from the simulations to show the mobility of D-loop (Fig. 5D). These lateral interactions stabilize the twisted conformation of the terminal subunits and reduce the solvent accessibility of the subunit P-1 D-loop by a factor of 5, which would compromise binding to the barbed end of an incoming actin subunit to elongate the filament. We confirm the prediction of Zsolnay et al. that subunits P and P-1 have bound ADP. The conformations of the terminal subunits in the simulations allows R39 and R62 of subunit P-1 to form novel networks of hydrogen bonds with subunit P (the sidechain of R39 with the OE1 of Q263 and the carbonyl oxygens of S271 and M269 of subunit P and with the carbonyl oxygens of its own R62 and L65, and R62 with subunit P E270). Our EM maps are consistent with these H-bonds, although the rotamers of the two arginine side chains are uncertain.

In agreement with previous work, we propose that the bent, twisted conformations of subunits P and P-1 compromise binding of an incoming actin monomer to the D-loop of subunit P-1 and the lateral side of subunit P for a substantial fraction of the time. This explains why subunit association is not a diffusion-limited reaction (Drenckhahn and Pollard 1986), but rather is limited by fluctuations of subunits P and P-1 between open and blocked states. The equilibrium distribution of the two states is not known, but judging from the nine-fold lower association rate constant for the association ATP-actin (Pollard 1986)), the pointed end appears to be available to bind ATP-actin monomers only 11% of the time and only 1% of the time in ADP-buffer, which favors the blocked, twisted state of the pointed end in MD simulations (Zsolnay, Katkar et al. 2020).

Both the P and P-1 subunits have wide open backdoors between subdomains 1 and 3 used for phosphate release and neither retain the γ-phosphate of ATP (Fig. 2ABC). The side chain of R177 is bent inward, opening the phosphate channel between it and N111. The slow rate of association of ATP-actin subunits provides ample time for ATP hydrolysis and the open backdoors allow the γ-phosphate to escape rapidly. Accordingly, all subunits in the pointed end reconstruction have bound ADP. The phosphate release gate is closed in more internal subunits including P-2 by a hydrogen bond between N111 and R177. The open gates of P and P-1 explain why the pointed end has a very low affinity for phosphate (Fujiwara, Vavylonis et al. 2007).

### Barbed end structure

The structures of barbed ends are similar in our 3.1 Å reconstruction from electron micrographs and molecular dynamics simulations (Zsolnay, Katkar et al. 2020). The terminal subunits are less twisted in our EM structure than in the simulations, but the subunits in both structures flatten progressively from the end (Fig. 6). The density for the terminal subunit in the EM map is much lower than the other subunits as anticipated by the MD simulations showing the D-loop as the only stable contact between subunit B and the other subunits, allowing the position of tethered B-subunit to fluctuate. The reconstructions show that subunits internal to subunit B hydrolyzed ATP and dissociated some of the γ-phosphate (subunits B-1 and B-2) or all of the γ-phosphate (internal subunits) during sample preparation. Based on the degree of flattening in MD simulations, Zsolnay et al. (Zsolnay, Katkar et al. 2020) predicted ATP hydrolysis is faster on subunit B-1 than subunit B, as observed in our reconstruction. Fast phosphate dissociation from barbed ends has not been measured directly but is expected from the observation that B-ends have a lower affinity for phosphate (Fujiwara, Vavylonis et al. 2007) and dissociate phosphate faster than internal subunits (Jégou, Niedermayer et al. 2011).

The overall agreement of the structures determined by MD simulations and cryo-EM supports the conclusion that binding site for incoming monomers on the barbed end of subunit B is readily accessible and that the tethered subunit B does not interfere with addition of subunit B+1. Once new subunit B+1 binds, lateral interactions with subunit B-1 will stabilize subunit B. The W-loop pocket of subunit B-1 is empty but open slightly wider than the internal subunits where M44 occupies the pocket, so subunit B-1 is ready to bind the D-loop of an incoming subunit. These properties are consistent with the association reaction being diffusion limited with a high probability (0.02) that a colliding monomer will bind (Drenckhahn and Pollard 1986).

Early studies of the barbed end did not have sufficient resolution to reveal details about the flattening of subunits or the status of their phosphate release channel. For example, Narita et al. (Narita, Takeda et al. 2006) docked a crystal structure of capping protein on the barbed end of 23 Å resolution cryo-EM structure of a capped filament.

A preprint by Oosterheert et al. (Oosterheert, F.E.C. et al. 2023) reports a 3.6 Å resolution structure of the barbed end of filaments prepared from Mg-ATP-actin monomers in the presence of DNase I, the FH2 domain of formin mDia1 and phalloidin. Their reconstruction lacks density for both the formin and the tethered terminal subunit that we call B. Oosterheert et al. may have been lost the terminal subunit and the FH2 domains, which have low densities in our maps, during local refinement in CryoSPARC with a soft mask around the terminal four subunits. Alternatively or in addition phalloidin binding at the interface of subunits B, B-1 and B-2 may have stabilized subunit B during more than 60 min before samples were frozen. We assume that their terminal subunit A_0_ is a stabilized version of our subunit B. The backdoor of their subunit A_0_ is partially open with 5 Å between the sidechains of N111 and R177 owing to disruption of a hydrogen bonding network including these two residues, H73 and G74. We measured an approximate distance of 5.3 Å between the sidechains of N111 and R177 in the weak density for subunit B and a reliable distance of 4.1 Å in subunit B-1. These spacings of N111 and R177 are much smaller than the wide open backdoors of subunit P and P-1 in our models. The spacing of the sidechains of N111 and R177 are 2.8 Å in Oosterheert’s subunit A_2_ and 3.6 Å or less in all internal subunits in our barbed end reconstruction.

### Extra densities associated with the barbed end

The extra densities on the sides of subunits B-1 and B-2 are too weak to build a model of the polypeptides de novo. These linear densities resemble superficially the bundle of helices of the FH2 domains of formin Cdc12, which was included with actin during polymerization. However, these densities are associated with SD1 rather than the barbed end groove and D-loops of adjacent subunits in the best available model from MD simulations of a barbed end with an FH2 dimer (from budding yeast Bni1) (Baker, Courtemanche et al. 2015). Additional work will be required to characterize these extra densities.

## Methods and Materials

### Actin purification

We purified actin from chicken breast muscle (MacLean-Fletcher and Pollard 1980). We extracted 4 g of muscle acetone powder with 80 mL of G-buffer (2 mM Tris-HCl, pH 8.0, 0.2 mM ATP, 0.5 mM DTT, 0.1 mM CaCl_2_, 1 mM NaN_3_) at 4°C for 30 min, pelleted insoluble materials and polymerized actin in the supernatant with 50 mM KCl and 2 mM MgCl_2_ at 4°C for 1 h. We dissociated tropomyosin from the actin filaments with 0.8 M KCl at 4°C for 30 min. After pelleting at 140,000× *g* at 4°C for 2 h, we dispersed the filaments with a Dounce homogenizer and then depolymerized them by dialysis against four changes of 1 L of G-buffer at 4°C over 3 days. After clarifying the depolymerized actin by centrifugation at 135,000× *g* at 4°C for 2 h, we applied the top 2/3 (∼10 mL) of the supernatant to Sephacryl S-300 gel filtration column equilibrated with G-buffer. This separates actin monomers (peak tail) from actin oligomers, capping protein and other minor contaminants (MacLean-Fletcher and Pollard 1980). We used an extinction coefficient of 1 OD_290_ = 38.5 μM to measure the concentration. Actin was stored at 4°C and used within two weeks.

### Cdc12 FH2 purification

We expressed recombinant *S. pombe* Cdc12 FH2 domains (residues 980 - 1420) fused to a tag containing thioredoxin domain and a hexa-histidine sequence in *E. coli* strain Rosetta 2(DE3) growing in LB medium. Expression was induced with 400 μM IPTG for 16 h at 22°C. Following lysis with sonication and lysozyme and pelleting insoluble materials, the protein was purified by affinity chromatography on a Ni-NTA column equilibrated with 50 mM Tris-Cl, pH7.5, 500 mM NaCl and 20 mM imidazole. FH2 domains were cleaved from the histidine tag by incubating for 12 h with PreScission protease at 4° C in 250 mM NaCl, 1 mM TCEP, 5 mM DTT, 20 mM Tris-Cl, pH 7.5 and washed from the beads with 500 mM NaCl, 50 mM Tris-Cl pH 7.5. The FH2 domains were concentrated by centrifugation at 3000 rpm with a 10k-cutoff membrane for 20 min and gel filtered on a Superdex 200 1.6/60 column in 100 mM KCl, 0.5 mM MgCl_2_, 0.5 mM EGTA, 5 mM Tris-Cl pH 7.5 and 0.5 mM DTT. The peak fractions were concentrated to 250 μM in 500 μL, frozen and stored at -80° C.

### Sample preparation for electron microscopy

We mixed 15 μL of 20 μM ATP-Mg-actin monomers with 15 μL of 6.0 μM Cdc12 FH2 domains in 100 mM KCl, 0.5 mM MgCl_2_, 0.5 mM EGTA, 5 mM Tris-HCl (pH 7.5), 15 mM HEPES (pH 7.0) and incubated for 140 min at room temperature to polymerize the actin. We used two type of grids, both glow discharged for 30 s at 25 mAmp, 36 mBar pressure. A sample of 3.0 μM of FH2 and 10.6 μM of actin monomers was applied to holey grids without a carbon film (Quantifoil Au 2/1 300 mesh from EMS). After diluting 4-fold, the same proteins were applied to holey grids with a 2 nm carbon film (Quantifoil Cu 2/2 200 mesh from EMS). Samples were frozen after a total of 145 min on the grids without carbon film and 150 min on the grids with carbon film. We used the following freezing parameters: blot force -12; wait time, 40 s; blot times, 3.0 s, with carbon film or 2.0 s without a carbon film; delay time, 2 s.

### EM image processing

We processed the images of short actin filaments using Relion 4.0 (Kimanius, Dong et al. 2021), including particle picking, motion correction and CTF estimation and map post-processing (Chou, Chatterjee et al. 2022). We wrote scripts called Fesp to identify short filaments with two ends and Fenda to align the ends (available on GitHub). Fesp displays both the coordinates (x, y) and the in-plane rotation (Fig. S1B). Fenda allows one to adjust the coordinates of each particle based on the coordinates of the class averages. We used two round of classification to align the ends of the particles (Fig. S1C). Comparisons with back projections of the two ends made from models of the middle of the filament were used to sort the class averages of the pointed and barbed ends (Fig. S1D-E). A 3D reconstruction from a sample of 448,222 pointed ends gave a map with 3.5 Å resolution and 471,000 barbed ends gave a map with 3.1 Å resolution (Table 1).

### Model building and refinement, and structural analysis and visualization

Our model of the ADP-P_i_-actin filament (PDB: 6DJN) fit well into the density for internal subunits of both reconstructions. We used Coot (Emsley, Lohkamp et al. 2010) to rebuild the structures of the terminal subunits of both structures. We used PHENIX (Adams, Afonine et al. 2010) to refine both structures in real space. We used PyMOL to calculate interdomain rotation (dihedral) angles and Chimera (Pettersen, Goddard et al. 2004) to calculate rise (subunit translation) and twist (subunit translation), RMSDs, and buried surface areas. Figures were generated using Chimera (Pettersen, Goddard et al. 2004) and Chimera-X (Pettersen, Goddard et al. 2021).

## Acknowledgements

Research reported in this publication was supported by National Institute of General Medical Sciences of the National Institutes of Health under award numbers R01GM026132 and R01GM026338. The content is solely the responsibility of the authors and does not necessarily represent the official views of the National Institutes of Health. The authors thank Shenping Wu, Jianfeng Lin, Marc Llaguno, Kaifeng Zhou, and Xinran Liu for managing the Yale Electron Cryo-Microscopy Resources, and the Yale Center for Research Computing for guidance and use of the research computing infrastructure

## Notes

### Competing Interest Statement

The authors have declared no competing interest.

