## Supplemental Figures for "Cryo-EM structures of both ends of the actin filament explain why the barbed end elongates faster than the pointed end"

Running Title: Actin filament ends

Supplemental materials

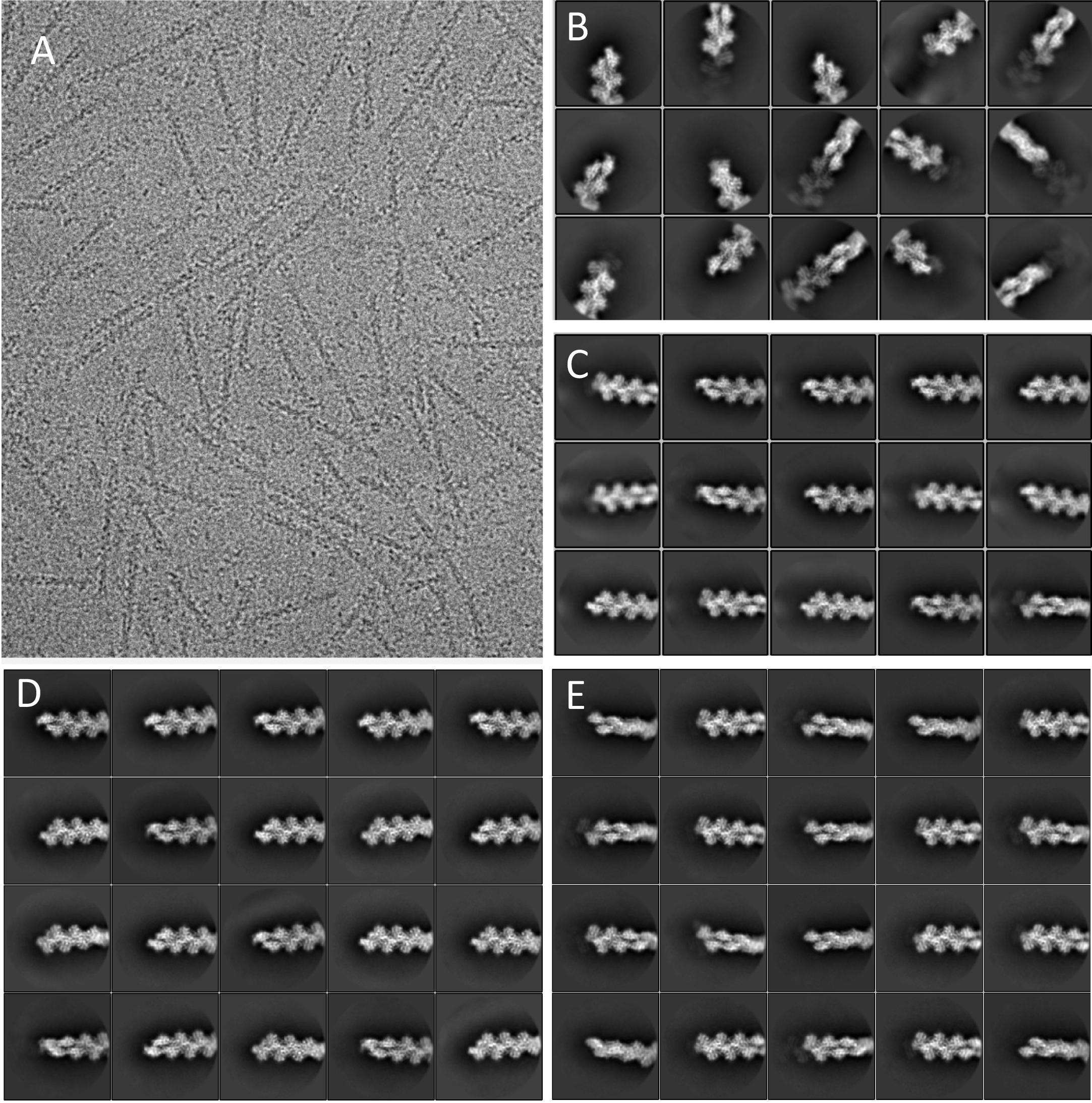

**Figure S1.** A, Electron micrograph of short actin filaments frozen on a Quantifoil grid with a carbon film. B, Class averages of filament ends identified by Fesp without correcting the orientation of each filament. C. Alignment of the ends of class averages with Fenda by moving each particle along the filament axis. (Note that due to the periodic features of the filament ends, moving a particle by one subunit also gives a high cross correlation value but can misalign the ends.) D-E. Manual separation of class averages into (D) pointed ends and (E) barbed ends by comparing with back projections of the two ends made from models of the middle of the filament.

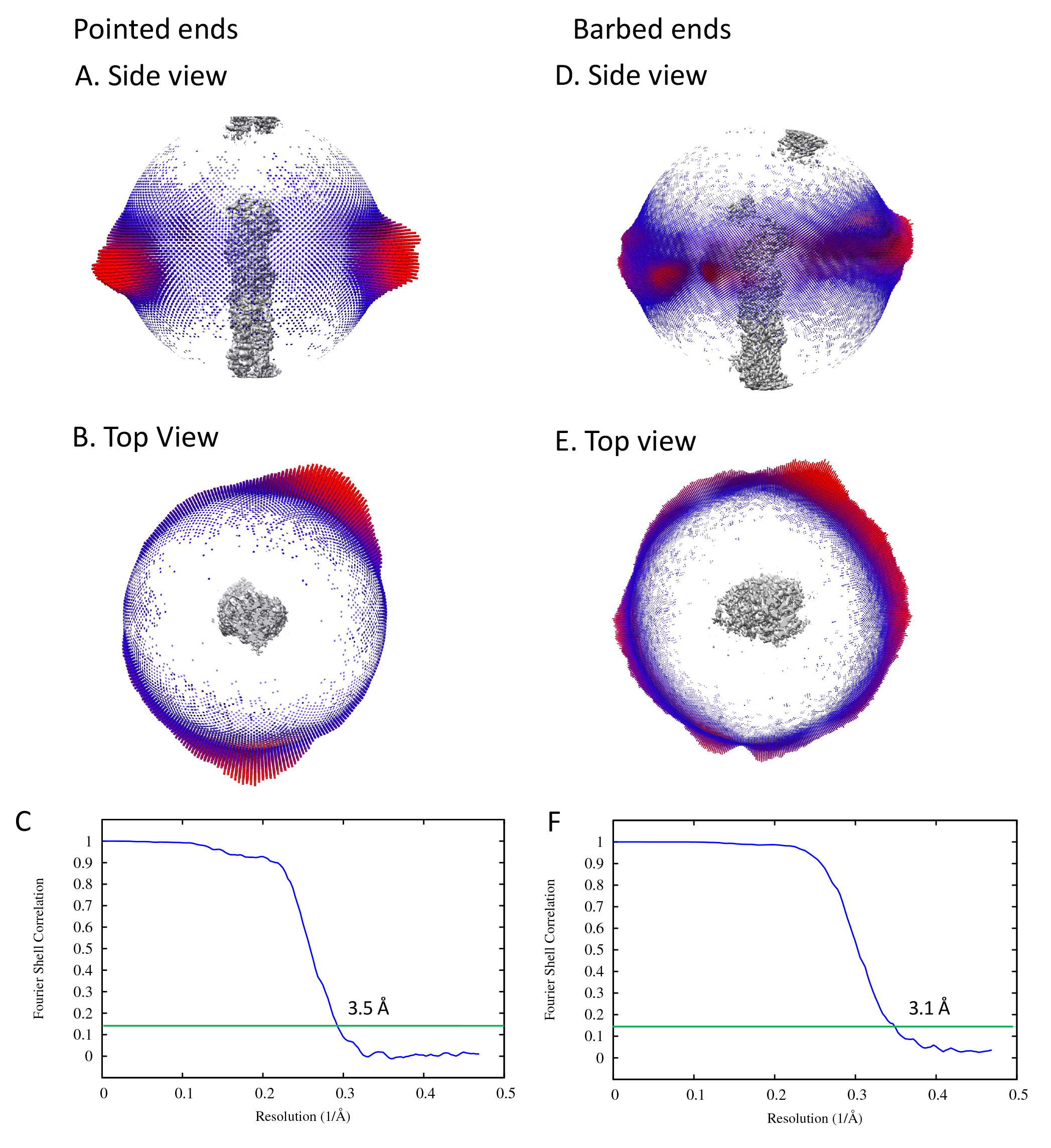

**Figure S2. Angular distributions of particles and resolution estimations.** A, B, D, E, Angular distributions of particles at the two ends, both shown as side views and top views. C, F, Resolutions of the reconstructions using Fourier Shell Correlation (blue curves) with the 0.143 criterion (green horizontal lines). A-C, Pointed end. D-F, Barbed end.
